# Conformational coupling by trans-phosphorylation in calcium calmodulin dependent kinase II

**DOI:** 10.1101/524660

**Authors:** Alessandro Pandini, Howard Schulman, Shahid Khan

## Abstract

**Abstract:** The calcium calmodulin dependent protein kinase II (CaMKII) is a dodecameric holoenzyme important for encoding memory. Its activation, triggered by binding of calcium calmodulin, persists autonomously after calmodulin dissociation. One (receiver) kinase captures and subsequently phosphorylates the regulatory domain peptide of a donor kinase forming a chained dimer as a first stage of autonomous activation. Protein dynamics simulations examined the conformational changes triggered by dimer formation and phosphorylation, aimed to provide a molecular rationale for human mutations that result in learning disabilities. Ensembles generated from X-ray crystal structures were characterized by network centrality and community analysis. Mutual information related collective motions to local fragment dynamics encoded with a structural alphabet. Implicit solvent tCONCOORD conformational ensembles revealed the dynamic architecture of Inactive kinase domains was co-opted in the activated dimer but the network hub shifted from the nucleotide binding cleft to the captured peptide. Explicit solvent molecular dynamics (MD) showed nucleotide and substrate binding determinants formed coupled nodes in long-range signal relays between regulatory peptides in the dimer. Strain in the extended captured peptide was balanced by reduced flexibility of the receiver kinase C-lobe core. The relays were organized around a hydrophobic patch between the captured peptide and a key binding helix. The human mutations aligned along the relays. Thus, these mutations could disrupt the allosteric network alternatively, or in addition, to altered binding affinities. Non-binding protein sectors distant from the binding sites mediated the allosteric signalling; providing possible targets for inhibitor design. Phosphorylation of the peptide modulated the dielectric of its binding pocket to strengthen the patch, non-binding sectors, domain interface and temporal correlations between parallel relays. These results provide the molecular details underlying the reported positive kinase cooperativity to enrich discussion on how autonomous activation by phosphorylation leads to long-term behavioural effects.

**Author Summary:** Protein kinases play central roles in intracellular signalling. Auto-phosphorylation by bound nucleotide typically precedes phosphate transfer to multiple substrates. Protein conformational changes are central to kinase function, altering binding affinities to change cellular location and shunt from one signal pathway to another. In the brain, the multi-subunit kinase, CaMKII is activated by calcium calmodulin upon calcium jumps produced by synaptic stimulation. Auto-transphosphorylation of a regulatory peptide enables the kinase to remain activated and mediate long-term behavioural effects after return to basal calcium levels. A database of mutated residues responsible for these effects is difficult to reconcile solely with impaired nucleotide or substrate binding. Therefore, we have computationally generated interaction networks to map the conformational plasticity of the kinase domains where most mutations localize. The network generated from the atomic structure of a phosphorylated dimer resolves protein sectors based on their collective motions. The sectors link nucleotide and substrate binding sites in self-reinforcing relays between regulatory peptides. The self-reinforcement is strengthened by phosphorylation consistent with the reported positive cooperativity of kinase activity with calcium-calmodulin concentration. The network gives a better match with the mutations and, in addition, reveals target sites for drug development.

## Introduction

The calcium calmodulin dependent protein kinase (CaMKII) is a multifunctional, multi-subunit eukaryotic protein kinase (EPK). It has key roles in calcium regulation of neuronal and cardiovascular physiology (1-3). EPKs mediate reactions whose malfunction promotes disease across a broad physiologic spectrum. The canonical EPK has a distinctive bi-lobed structure that exploits diverse strategies to achieve allosteric regulation (4).

CaMKII has a canonical kinase domain (KD) tethered via a linker to an equally well-conserved association domain (AD) that forms a central hub of the dodecameric holoenzyme with two hexamer rings that stack with mirror symmetry. Linker diversity generates isoforms and splice variants. The kinase is activated by rises in cellular calcium that enable calcium-calmodulin (Ca^2+^/CaM) to bind to and displace an autoinhibitory regulatory domain. The autoinhibitory domain occludes interaction of CaMKII with anchoring proteins, such as NMDA receptor subunit GluN2B (5, 6). More subunits are activated at increasing frequency of calcium pulse trains. The tuning frequency depends on the switching kinetics between the holoenzyme open and closed states (7).

In the closed state, the substrate binding surface is occluded by an auto-inhibitory segment composed of a N-terminal segment (R1) and C-terminal α-helices (R2, R3) and ATP affinity is low (8). R1 contains the primary auto-phosphorylation site (T286 in α subunit, T287 in others), as well as residues for O-GlcNAC modification (S279) and oxidation (M280, M281 (δ isoform)). R2 has a CaM recognition motif. Ca^2+^/CaM binding displaces the regulatory segment to switch CaMKII to the open state, followed by segment capture and T286 autophosphorylation by an adjacent, open (activated) subunit (8, 9) in the holoenzyme. R3 has additional threonine residues (T305, T306) that are primarily phosphorylated when CaM dissociates from a Ca^2+^-independent (autonomous) kinase (10, 11) to counteract T286 phosphorylation (12). X-ray crystal structures show that R1 and its binding determinants in the KD core adopt different conformations. Electron spin resonance (ESR) reports that R1 becomes unstructured upon T286 phosphorylation, while R2 and R3 are disordered in solution (13).

Persistent CaMKII activity autonomous of Ca^2+^/CaM underlies conversion of transient synaptic stimulation to long-term potentiation (LTP); a fundamental problem of neuronal CaMKII biochemistry that underlies certain forms of learning and memory (6, 14-17). Recent studies have identified more than a dozen human mutations in CaMKII that result in various degrees of learning disabilities (18, 19). These mutations when mapped onto structure localize largely with R1 or mutations that alter KD interactions with substrates rather than R2. The previously described binding mutations form S (substrate binding) and T (Thr286 docking) sites (20, 21)). The substrate binding determinants overlap with the R1 binding groove that can be partitioned into A, B and C sites (8, 22). The T-site is eliminated upon release of the autoinhibitory segment by rotation of helix αD whose movement helps form the B site (23). An outstanding issue is whether the human mutations act independently or as part of a collective network to disrupt allosteric communication responsible for frequency tuning (24). The conventional form of autonomous activity follows T286 trans auto-phosphorylation, which maintains activity after dissociation of Ca^2+^/CaM. Structures of CaMKII complexes with R1 regulatory segments of one subunit (“donor”) captured by the adjacent subunit (“receiver”) upon Ca^2+^/CaM binding, give important snapshots into trans-phosphorylation (8, 9). A hydrophobic clamp of helix αD residues within the receiver KD captures the R1 in an interaction that propagates throughout the crystal lattice.

Here, we used tCONCOORD, based on stochastic distance constraints (25), and explicit solvent molecular dynamics (MD) as complementary methods to address the mechanistic basis underlying the medical phenotypes of the human mutations. tCONCOORD ensembles from KD structures described collective motions and conformational coupling between the nucleotide and substrate binding sites. MD examined fine-grained architectural modulations due to primary site phosphorylation on the long-range allosteric network to establish an analytical framework for the coupling within and between subunits. We characterized the long-range allosteric relays in a KD dimer extracted in silico from the crystal lattice as well as KD monomer structures. Two strategies analyzed the network architecture based on mutual information. First, the centrality profile of the complete network of local correlated motions was determined to identify relays of the most strongly coupled fragments. Second, the protein was partitioned into contiguous sectors, with coupled dynamics, that act as semi-rigid bodies – henceforth defined as “communities” (26). Community size and cross-talk tracked the transition between the auto-inhibited and phosphorylated states.

We show that the captured R1 acts as a strained spring, rather than a “grappling hook” (8) to couple nucleotide and substrate binding sites (27, 28). The capture freezes out C-lobe core motions to generate long-distance signal relays between R1 segments of adjacent subunits. The human mutations align along the relays. T286 phosphorylation strengthens the relays to suggest how it might facilitate inter-subunit conformational spread and modulate affinities at distant sites for calmodulin (29) and cytoskeletal actin (30). Notably, we identify a hitherto uncharacterized sector to guide the design of allosteric inhibitors. The simulations should complement experimental ESR (13) and FRET measurements (31, 32) on activated CaMKII dimers for dissection of the activation mechanism.

## Results

### 1. The human mutations localize to the KD C-lobe and R1

Figure 1A provides a road-map of the architectural elements investigated in this study. The R1 with T286 binds to a groove within the adjacent receiver KD C-lobe that overlaps with the receiver’s S-site and has been sub-divided into three pockets A, B and C (8) in the nematode *C.elegans* KD dimer (PDB ID:3KK8)). The A-site forms a canonical substrate docking interaction with the ATP binding cleft that contains the DFG motif (D156FG158); the B-site forms a hydrophobic patch with helix αD; the C-site has acidic residues for salt-bridges with R1. Residue positions with mutations in the human homolog that result in learning disabilities as well as S-site and T-site mutations are mapped onto the donor KD. The human mutations localized to R1 (I272, H282, R284, T286), or overlapped with, or were adjacent to the S and T-sites (F98S, E109, P138). However, a significant fraction mapped elsewhere.

**Figure 1:**
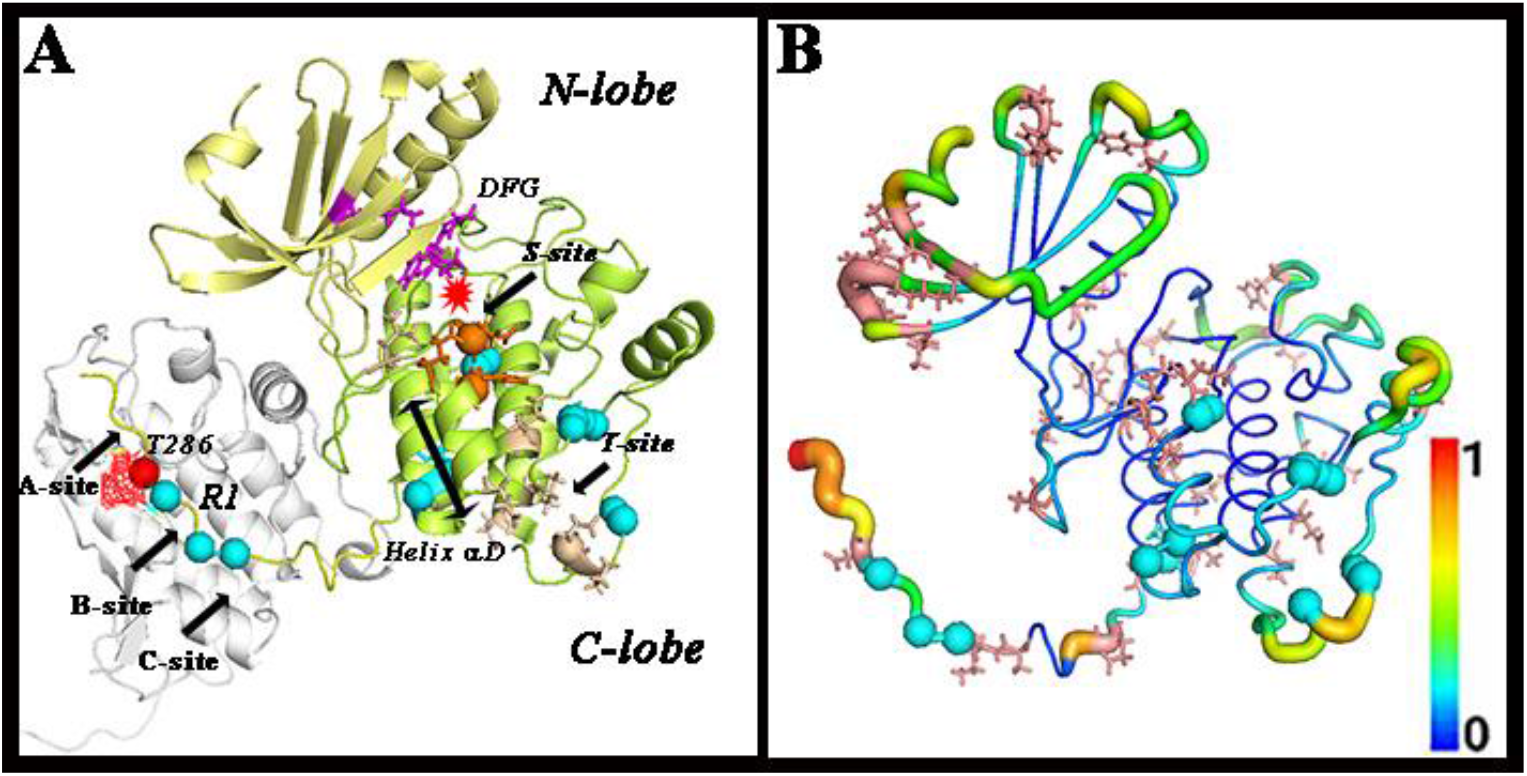
CaMKII Functional Mutations. Spheres (cyan) show residue positions where mutations cause learning disabilities in humans. **A. CaMKII KD architecture.** Structure of the CaMKII kinase domain (3KK8) with N-terminal (pale yellow) and C-terminal lemon) lobes and R1 regulatory segment (yellow). Spheres mark S (orange) and T site (yellow) binding mutations (20), primary auto-phosphorylation site (T286 (red with mesh)). Helix αD (double arrow bar). R1 is bound to a groove in an adjacent subunit (white)). The groove has three sites – A, B (and C (8). Asterisk (red) marks magnesium ion location. **B. CaMKII KD flexibility and sequence variation.** Room mean square fluctuation (RMSF) profile computed from the 3KK8 ensemble. Stick (salmon red) representations identify variable residue positions as identified from the MSA (Figure S1). Vertical bar indicates color scale (1 = high (red); 2 = low (blue)).

The flexibility profile recorded by the root mean square fluctuations (RMSF) of residue positions computed from the tCONCOORD ensemble of (Figure 1B) is consistent with the b-factor values in the crystal structure. Variable residue positions, identified from the multiple sequence alignment (MSA) of > 500 KD sequences, localized to surface accessible loops when mapped onto 3KK8. The MSA shows strong sequence conservation between species and minimal differences between isoforms within species. The phylogenetic tree indicates that nematode and human sequences are among the most distantly-related (**Supporting Information Figure S1**). Secondary structure predicted from the MSA for the *C. elegans*, rat and human sequences matched that observed in the crystal structure. The predictions back ESR evidence for R2 and R3 disorder (13). The human mutations localized to mainly conserved residue positions in both rigid and flexible segments of the KD C-lobe and R1.

### 2. The captured R1 forms the central node of the allosteric network

We analyzed local protein dynamics to decipher allosteric signal propagation Three, separate, explicit solvent MD replica runs of the 1.7 angstrom resolution structure of the phosphorylated 3KK8 dimer and its non-phosphorylated derivative were performed. The 3KK8 dimer was extracted from the crystal structure electron density with the R1 of the donor KD captured (“bound”) by the adjacent receiver; while the R1 of the receiver KD was unconstrained (“free”), immersed in solvent. Four-residue fragments were encoded by the structural alphabet (SA) (33) in structures within the conformational ensembles and trajectories. In the network, the nodes were the fragments (n = 560), while edges were the correlations (n*(n-1)/2) = 156520). The profile reflected the contribution of individual fragments to the dynamics driving the PC motions and is optimal for detection of network nodes (Figure 2A). The T and S sites emerged as prominent nodes in addition to the captured R1 However, the network architecture is too dense to visualize the spatial extent or interaction strength of the connections formed by individual nodes.

**Figure 2:**
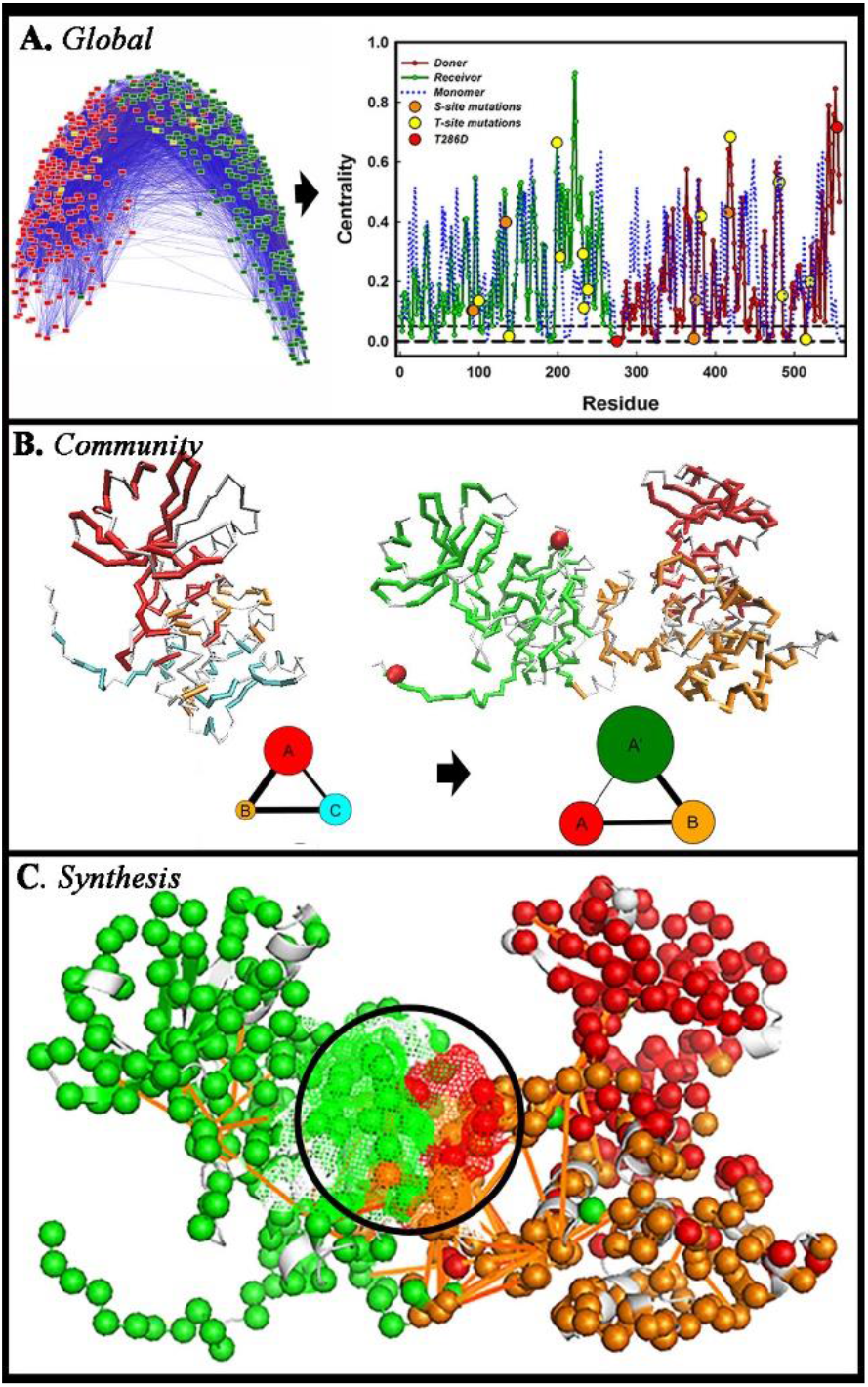
The captured R1 forms the community hub of the chained dimer. **A.** The global network (donor KD (red), receiver KD (green) and its eigenvector centrality. Spheres mark S (orange) and T site (yellow) mutant residue positions and TPO286 (red). **B.** Community membership of the CaMKII KD (KK8) monomer (open-form) and dimer (ribbon representations) mapped onto the structures with associated community graphs (node diameter = community size; edge thickness = coupling strength). **C.** Distinct communities (color-coded C^α^ atoms (spheres) and top fragment-fragment couplings (nMI > 0.2 (orange lines)) are superimposed onto the structure. Mesh (circled) marks the captured R1 and contacting residues.

Communities (n > 3), each represented as a node, greatly simplify network visualization of the connectivity by encoding concerted domain motions intermediate between global motions and residue-level RMSF fluctuations. Community membership and interaction strength were computed based on spectral decomposition (34, 35). Our methodology reproduced the published community map of the PKA catalytic subunit (PDB ID: 1CMK) (26) and showed that the CaMKII KD had similar dynamic architecture to PKA with the ATP binding site forming the community hub (Figure 2B, **Supporting Information Figure S2**). Three major communities (A, B, C) converged at the ATP site. The N-lobe community A included the ATP binding cleft and part of the A-site. The C-lobe was split along its centre into community B with B-site helix αD and community C with the C-site acid residue pocket. Donor KD communities B and C coalesced with dimer formation (B’), while one community (A’) accounted for the receiver KD. The top couplings from the global network scored on mutual information (nMI) were superimposed on the community architecture (Figure 2C). The captured R1 replaced the ATP binding site as community hub. The couplings congregated around it to link the donor A site with the receiver B and C sites.

### 3. The DFG motif and helix αD link communities in diverse KD structural states

We compared the 3KK8 KD with other CaMKII KD structural states in the Protein Data Bank (PDB) to first correlate community dynamics with monitors of kinase activity. Activation involves inward motion of the DFG side-chains for nucleotide binding and inward tilt of helix αD for contact with substrate peptides in many diverse EPKs (36). The states could be grouped into three categories - open, inhibitor-bound and auto-inhibited. tCONCOORD conformational ensembles were generated. The dominant conformations for each ensemble were identified from cluster analysis and aligned (Figure 3A). The two open activated states (3KK8, 2WEL) and the inhibitor-bound form (3KL8) had inwardly oriented DFG motifs and αD helices. In contrast, the auto-inhibited states (2BDW, 2VN9, 3SOA) had outwardly rotated helix αDs. While 3SOA has an outwardly oriented DFG motif, the loops for 2BDW and 3SOA had inward orientation albeit at a position distinct from that assumed by activated states.

**Figure 3:**
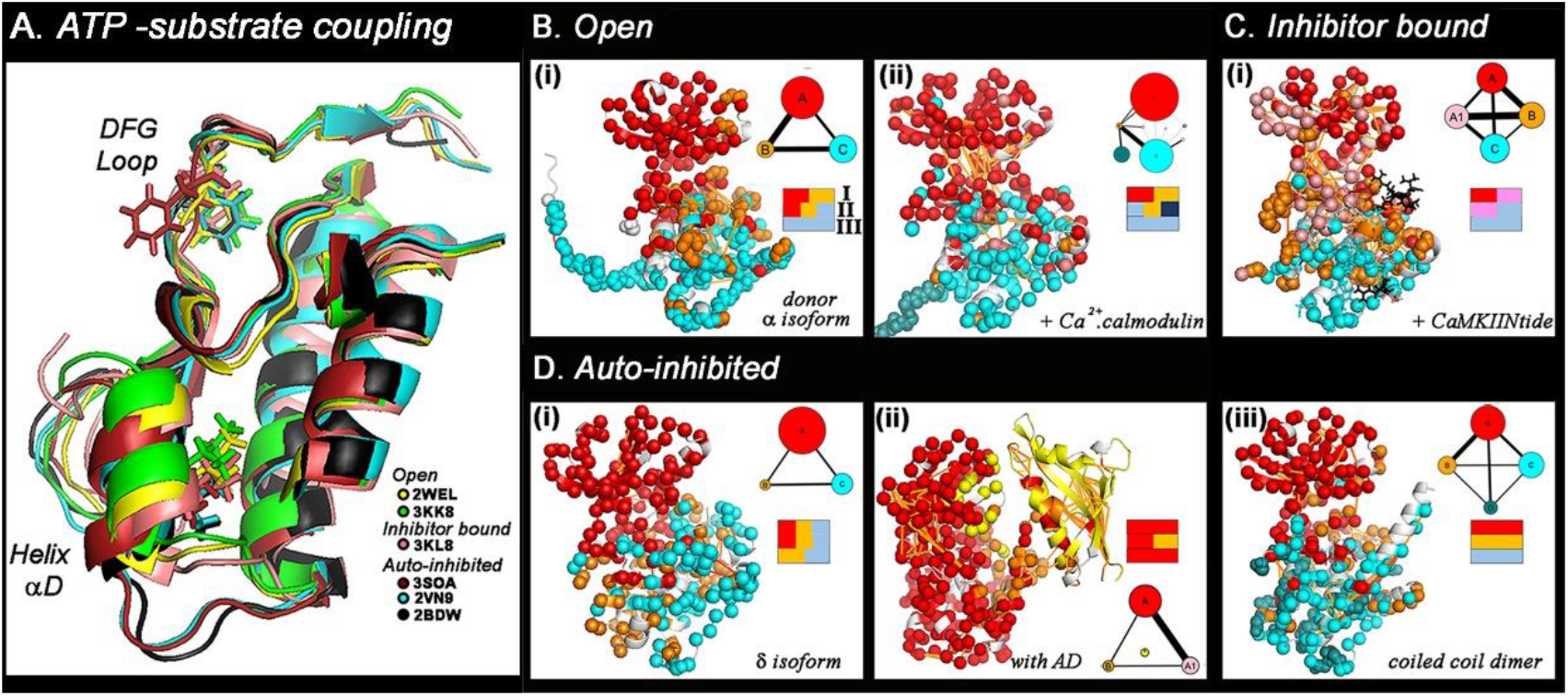
A. Analysis of the coupling between ATP and substrate binding sites. Aligned KD C-lobe crystal structures. RMSD’s (angstroms) - 2BDW (reference), 2VN9 (0.30), 2WEL (0.34), 3KK8 (0.23), 3KL8 (0.62), 3SOA (0.29). **Community dynamics in KD crystal structures**. Colors (red (N-lobe), orange (central C-lobe), cyan (basal C-lobe)) denote major communities following the 3KK8 donor KD community structure (Figure 2B). Top dynamic couplings (orange lines) are mapped on the structures as in Figure 3C. **Insets**: Community membership of the ATP binding motif D156FG158 **(I)**, helix αD L_97_FEDIVAR_104_ **(II)** and C site P_235_EWD_238_ **(III)** in **B - Open forms. (i)**. Nematode CaMKII with free R1 (3KK8 donor KD), **(ii)**. human δ with bound calmodulin (2WEL). The segment bound to calmodulin forms a distinct community (navy) with community B partitioned into smaller communities. **C - Inhibitor-bound form. (i)** Nematode CaMKII with bound CaMKIINtide (3KL8). **D - Auto-inhibited forms. (i)**. Human δ isoform (2VN9). **(ii)**. Complete (KD with AD) human CaMKII α (3SOA). Contacts between the N-lobe and AD form a distinct community (yellow). **(iii)**. Nematode CaMKII stabilized by R2 helix (coiled-coil dimer) (2BDW).

We next catalogued the community membership of the DFG motif (**I**), fragment L97FEDIVAR104 from B-site helix αD (**II**) and fragment P235EWD238 from the C-site acid pocket (**III**) (Figure 3B-D). The open forms (Figure 3B) differed in their community architecture owing to the bound calmodulin. The calmodulin fragmented community B, with R1 dynamics uncoupled from the C-lobe. In both forms, fragments I and II had multiple membership in communities A and B. The CaMKIINtide inhibitor peptide associated with sites A, B and C similarly to the captured R1 (8) (Figure 4C). Accordingly, community interactions are substantially increased, even though the communities did not coalesce. Community interactions were weak in the auto-inhibited states (human δ, human α full-length subunit, and a KD extracted from the nematode coiled-coil dimer) (Figure 4D). Community A extended into the C-lobe when AD contacts limited the relative motions of the N and C-lobes to each other (3SOA). Fragment II connected the dominant communities as did fragment I in the auto-inhibited human δ, as well as the human α KD.

**Figure 4:**
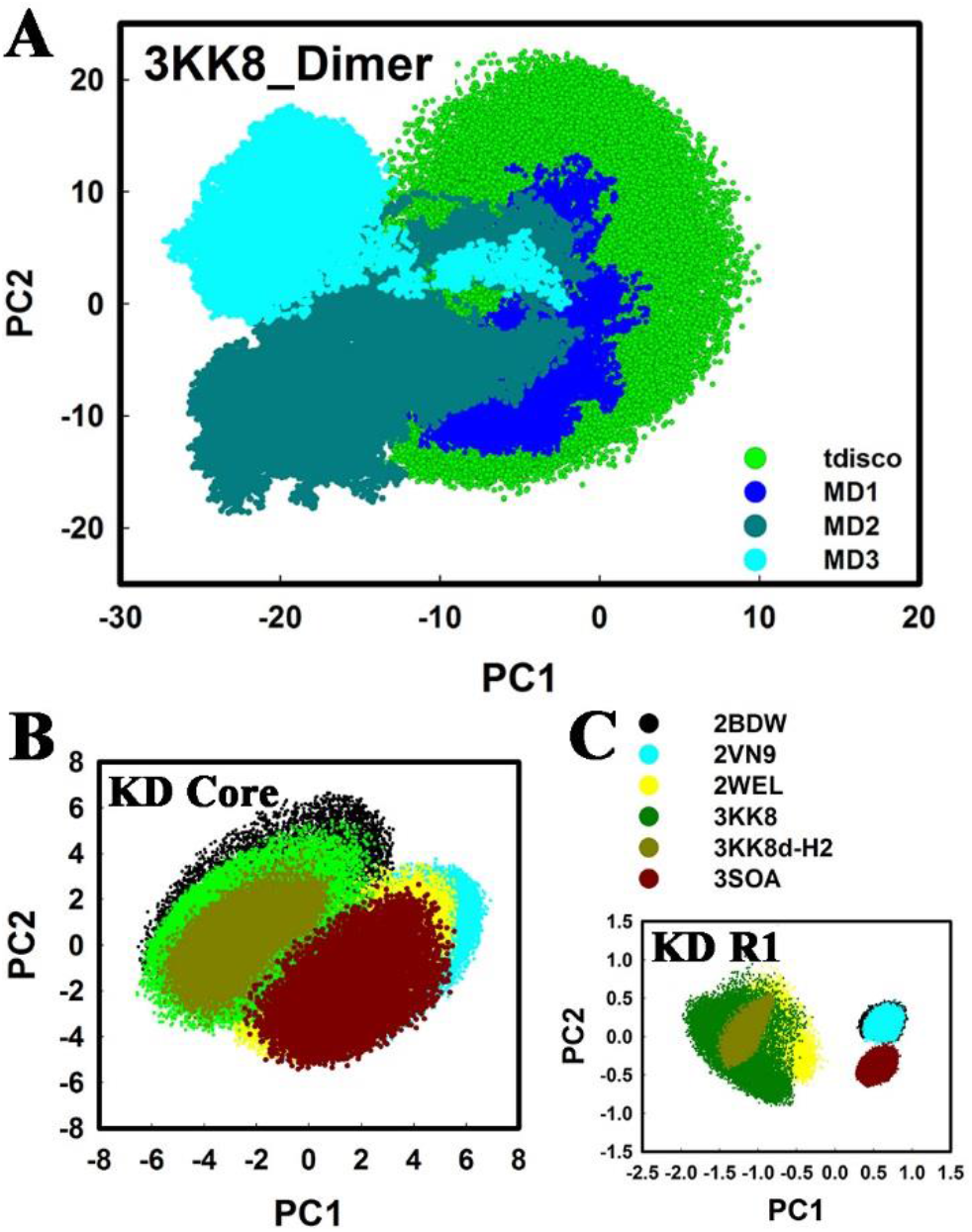
The captured R1 inhibits collective motions of the kinase core. PC1PC2 plots show the relation between the collective motions recorded by the amplitudes of the principal components PC1 and PC2. **A.** PCiPC2 plot for the 3KK8d dimer. **B, C.** PC1PC2 plots of the (**B**) KD core (N-lobe + C-lobe minus R1 and (**C**) R1 of the donor (3KK8) and receiver (3KK8-H2) KDs of the 3KK8 dimer compared with plots for the KD from 4 crystal structures (auto-inhibited (2BDW) and CaM bound (2WEL) C. elegans; autoinhibited human δ (2VN9) and α (3SOA) isoforms. Structure color code follows 3A.

In conclusion, the coupled ATP binding DFG motif (Fragment I) and helix αD (Fragment II) form a universal hub in activated and inactivated CaMKII KD states based on their multiple community membership. The hub is robust to variations in community interaction strength. Inward movement of helix αD is a diagnostic for activation, but that of the DFG motif not necessarily so

### 4. The captured R1 freezes KD core motions

We next characterized collective motions, extracted by principal component analysis (PCA), to better understand the monomer to dimer transition. Principal motions (PCs) primarily delineated relative motions of N and C lobes. The amplitude of the first two PCs accounted for >70% of the combined amplitude. PC1PC2 plots compared the amplitudes of the dimer motions obtained by MD and tCONCOORD (Figure 4A). The 300 ns MD runs reached steady-state configurations between 50 to 150 ns. The PC1PC2 conformational space sampled during each run was typically > 2/3 the space sampled by the tCONCOORD ensemble. The space sampled by the combined MD trajectories was comparable (> 90%) to the latter ensemble, with considerable overlap between the two.

Motions of the subunit KD cores and R1 segments were compared to evaluate the extent to which captured R1 motions determined dimer flexibility. The mechanical hinge and shear surfaces responsible for the PC motions formed community interfaces, with the ATP binding cleft as hub in the monomer (**Supporting Information Movie S1**). The captured R1 was revealed to be the central hinge or hub for the dimer PC motions (**Supporting Information Movie S2**). The PCA shows that the captured R1 acts as an allosteric effector to freeze-out the motions of the kinase core.

Simulations on other CaMKII KD structures revealed the principal dimer motions were threefold greater relative to monomer KD core motions. PC1PC2 spread of the receiver KD core was reduced (> 60%) relative to the spread of donor KD core and, indeed, KD cores from the other structures (Figure 4B). The PC1PC2 plots for the R1 segment ensembles formed two distinct groups (Figure 4C). The group with smaller spread represented auto-inhibited structures. The cluster with larger spread (4x) consisted of the captured R1 and free R1 segments. The PC1PC2 spread of the captured R1 was reduced 50% relative to the free R1 segments. The scenario most simply consistent with these results is of a 3-state transition between autoinhibition and activation. R1 is immobile docked as pseudo-substrate (“auto-inhibited” state), mobile when free in solution (“open” state) and constrained upon subsequent capture (“activated” state). The mobility of the KD core does not change when R1 is displaced but it is notably reduced upon R1 capture. Therefore, R1 freezes KD core collective motions more effectively when captured than when docked as pseudo-substrate, but with compensatory increase in its own flexibility.

### 5. Energetics of the chained dimer

Allosteric signal transduction has both enthalpic (Δ*H*) and entropic terms (Δ*S*) that contribute to the free energy change (Δ*G*).

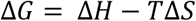

Enthalpic changes result from the free energy of solvation and residue pK (log_*e*_ *K*) shifts due to modulation of the dielectric constant upon complex formation or covalent modification.

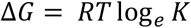

The normalized mutual information (*nMI*) provides an information theoretic measure of the coupling. This is consistent to the Shannon entropy *H*(*X*) for pairs of fragments albeit incomplete because of finite sampling (37). For any variable (*X*), the entropy Δ*S*(*X*) is related to the number of microstates and their probability, where *k_B_* Is the Boltzmann constant.

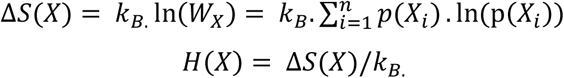

We compared the end-to-end distance distributions of the captured relative to the free R1 to assess strain (Figure 5A, **Supporting Movies S3-S4**). The captured R1 was constrained to a highly-extended subset of conformations. Secondary structure analysis based on the SA (Figure 5B) revealed the extended configuration resulted from melting of two R1 α-helical segments N551RERV and D563VDCL; while the intervening alanine-rich V555ASAI sequence largely retained α-helical character. The result was in line with mean residue helical propensity (38) of the fragments (−0.18 (V555ASAI) > - 0. 29 (N551RERV) and 0.49 (D563VDCL) kcal / mol).

**Figure 5:**
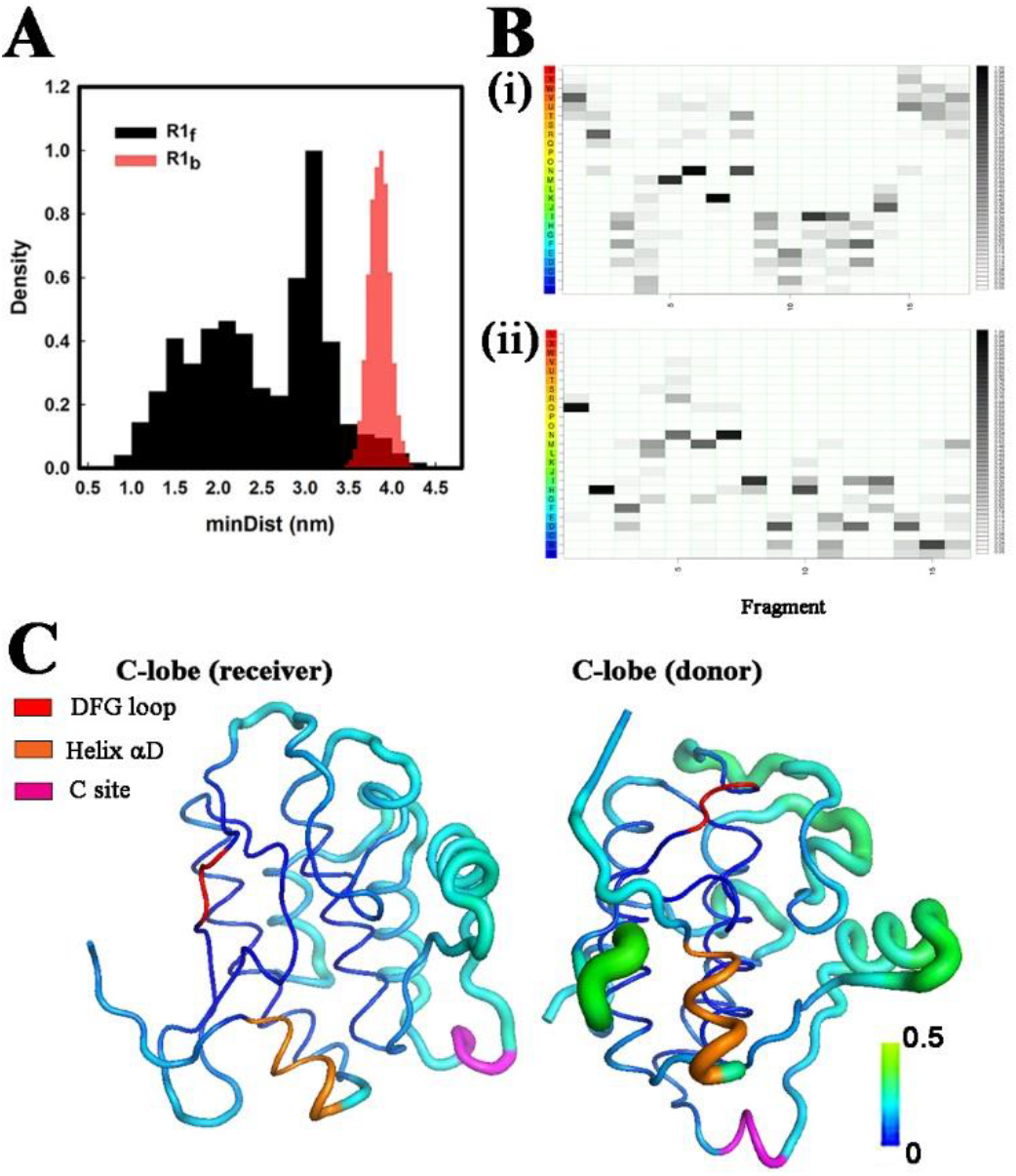
Capture strains R1 and freezes the receiver C-lobe. **A.** End-to-end distance distributions of the free (black) and bound (red) R1 segments. **B.** SA distribution profile showing loss of α-helical character upon transition from (**i**) free to (**ii**) bound. The SA letters are color coded (red = α -helical -> blue = β sheet). **C.** RMSF fluctuations from one MD replica run (TPO form) mapped onto the 3KK8 dimer. R1 capture freezes out helix αD L_97_FEDIVAR_104_ (orange) and interface motions in the C-lobe of the receiver core. D_156_FG_158_ motif (red), C-site P_235_EWD_238_ (magenta). Vertical bar indicates RMSF value on the same color scale shown in Figure 1B.

We focussed on the C-lobe to evaluate the compensatory decrease in flexibility of the core with greater precision (Figure 5C). Comparison of the RMSF profile of the dimer C-lobes shows that reduced flexibility of the B site helix αD and the interfacial surface of the receiver lobe are responsible for the overall decrease reported by PCA (Figure 4B).

### 6. Electrostatics of T286 phosphorylation

The solvent exposed area and electrostatic properties of the phosphorylation site were calculated for the most populated conformations extracted by cluster analysis. Dimer conformations from each set of MD replicas, as well in the tCONCOORD ensemble with the phosphomimic mutation (T286D) (1) were analysed. Dimer contacts were localized to the C-lobes. Contact between R1 of the donor subunit with the receiver subunit core upon capture accounted for much of the stabilization due to solvation. The total buried surface area due to subunit capture was 2319±42 A^2^. The contribution to this value due to R1 capture was 1765±81 A^2^ (~ 75%). The free-energy difference due to the decrease from solvation was twice as great for the phosphorylated (−7.4 kcal/mole) versus the non-phosphorylated (−4.2 kcal/mole) form.

Computed Poisson-Boltzmann electrostatic fields are shown in (Figure 6A). The R1 binding pocket was more acidic and the dimer interface more polar when T286 was phosphorylated (TPO286) or mutated (D286). Polar residues with large (> 0.5) pK shifts between the T286 and TPO286 forms were identified by comparison of the dominant dimer configurations (Figure 6B). These residues localized to the C-lobes around the R1 binding pocket and the subunit interface (Figure 6C). Phosphorylation shifted the surface of the R1 binding pocket to acidic values, while its interior became more hydrophobic. The measured pK shifts accounted for the more rigid interface.

**Figure 6:**
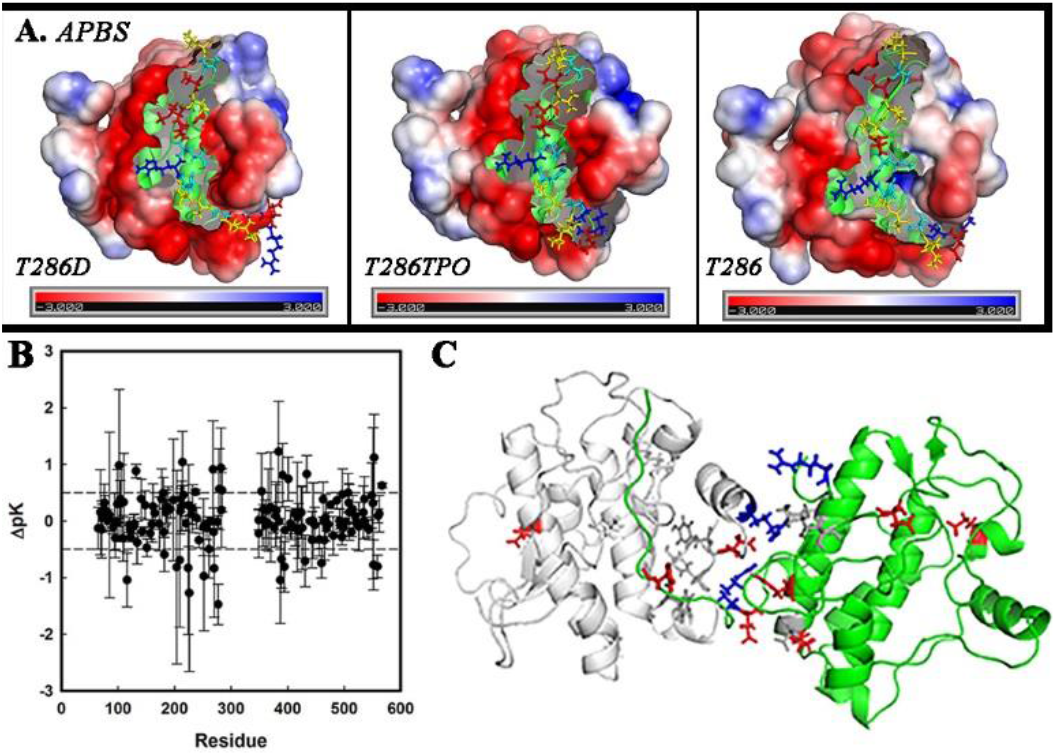
Primary site phosphorylation increases R1 cross-talk. **A.** Changes in the electrostatic environment (red = acidic, blue = basic, white = hydrophobic) triggered by T286 modification (-> TPO phosphorylation or -> *D* mutation). The binding crevice becomes more acidic while the adjacent subunit interface becomes more basic. Color scale is −1 to + 1*^k_B_T^/_e_ mV*. **B.** Phosphorylation dependent residue pK shifts (mean + standard error). **C.** Map of residue (stick side-chains) pK shifts > 0.5 onto the C-lobes of the 3KK8D.pdb dimer. Colors denote residues shifted towards acidic (red), basic (blue) or more neutral (dark grey) values. The computed free energy change is −1.1 kcal / mole.

### 7. The allosteric network forms a self-reinforcing R1 relay

Community membership mapped onto structure revealed the captured R1 segment and its binding pocket formed a central community (B’) distinct from A’. B’ had strong interactions with donor KD communities A and B. The adjacent R1 segments were linked by long-range couplings between B’ and B. The nucleotide binding pocket and adjacent fragments of its donor KD also linked to the captured R1 (Figure 7A). Superposition of the network centrality profile onto the structure delineated a signal relay from the donor A-site to the captured R1 via helix αD (B-site) in the receiver to its R1 (Figure 7B, **Supporting Movie S5**).

**Figure 7:**
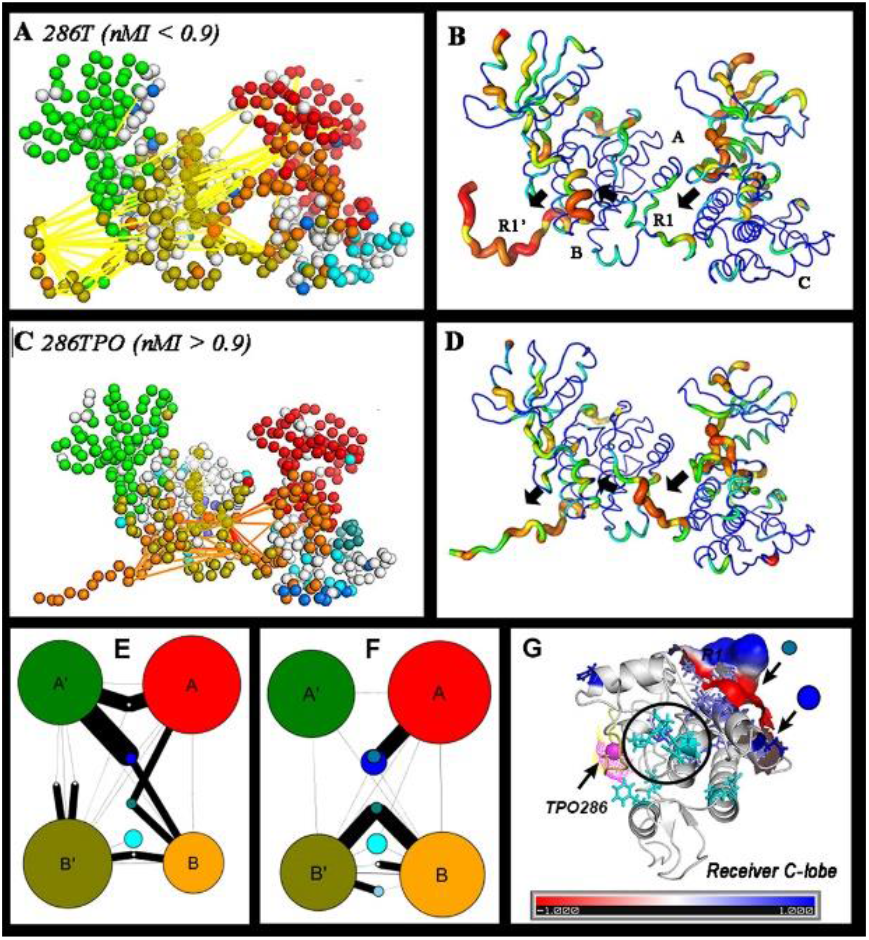
T286phosphorylation increases R1 cross-talk. **A.** Community membership and top couplings (yellow lines) of the native dimer. **B.** Eigen-centrality map of the native dimer. **C.** Community membership and top couplings (orange lines) of the phosphorylated (TPO) dimer. **D.** Eigen-centrality map of the TPO dimer. G -> R color code (bar) and thin -> fat backbones indicate increasing values. Arrows (black) indicate shortest pathway linking the substrate subunit A site and R1 with the enzyme subunit B site and R1. **E, F.** Community graphs for the native (E) and TPO (F) dimer show parallel signal processing between the donor R1 (in community B) and the receiver R1 (in community B’). **G.** Surface protrusion and electrostatics (basic polar) of the receiver KD protein sector (249D-259R, 263D) that forms the intermediate community (teal sidechains) mediating the dominant B -> B’ signal relay in the TPO286 dimer. The more distributed signal community mediating the A -> B’ signal relay (blue sidechains) increases size by recruiting the same sector from the donor KD upon phosphorylation. Three proline residues (P212, P213, P235) (circled) targeted by human mutations (cyan sidechains) cluster between the captured TPO286 (magenta sphere) and the teal community. R1 (yellow).

R1-R1 communication was increased upon phosphorylation (T -> TPO) due to the creation of strong couplings (nMI > 0.9) between captured R1, the donor A and receiver B and C sites (Figure 7C). The dominant signal relays were remarkably similar to those representing the top couplings from the tCONCOORD D286 ensemble. The map of TPO dimer centrality profile highlighted the relay was more localized to the captured R1, donor A, C and receiver B sites (Figure 7D).

Parallel network signal processing in the networks was quantified with community size (residue number) and interaction strength represented, as in Figure 2B, by community graphs (**Figures 7E, F**). Phosphorylation strengthen B -> B’ interactions at the expense of A’ interactions. Two small communities mediated A -> B’ and B -> B’ signal relays. These were generated upon R1 capture and strengthened upon phosphorylation. A sector common to both communities comprised a loop between C-lobe α-helices 9 and 10 that is well conserved, polar in character and at the opposite face to the R1 binding site (Figure 7G, **Supporting Movie S6**). The loop is an attractive target for the design of allosteric inhibitors based on these characteristics.

### 8. Temporal correlations within the R1 relay

Transmission of information between subunits encoded by the phosphorylation-dependent couplings was analysed to understand signal relay kinetics. The core network of long-range allosteric couplings (n = 115, nMI > 0.09) linked the bound and free R1 to sites A, B and C (Figure 8A). A patch of three hydrophobic residues that attach R1 to B site helix αD (I101, A279”, I280”) formed the central node. This node is less compact in the native (T286) relative to the phosphorylated (TPO286) dimer. Most human mutations causing learning disabilities map within or close to the central and accessory network nodes. In addition to communication between the intermediate relay communities and R1 (Figure 7G), the proline (P212, P213, P235) mutations could also regulate interface flexibility.

**Figure 8.**
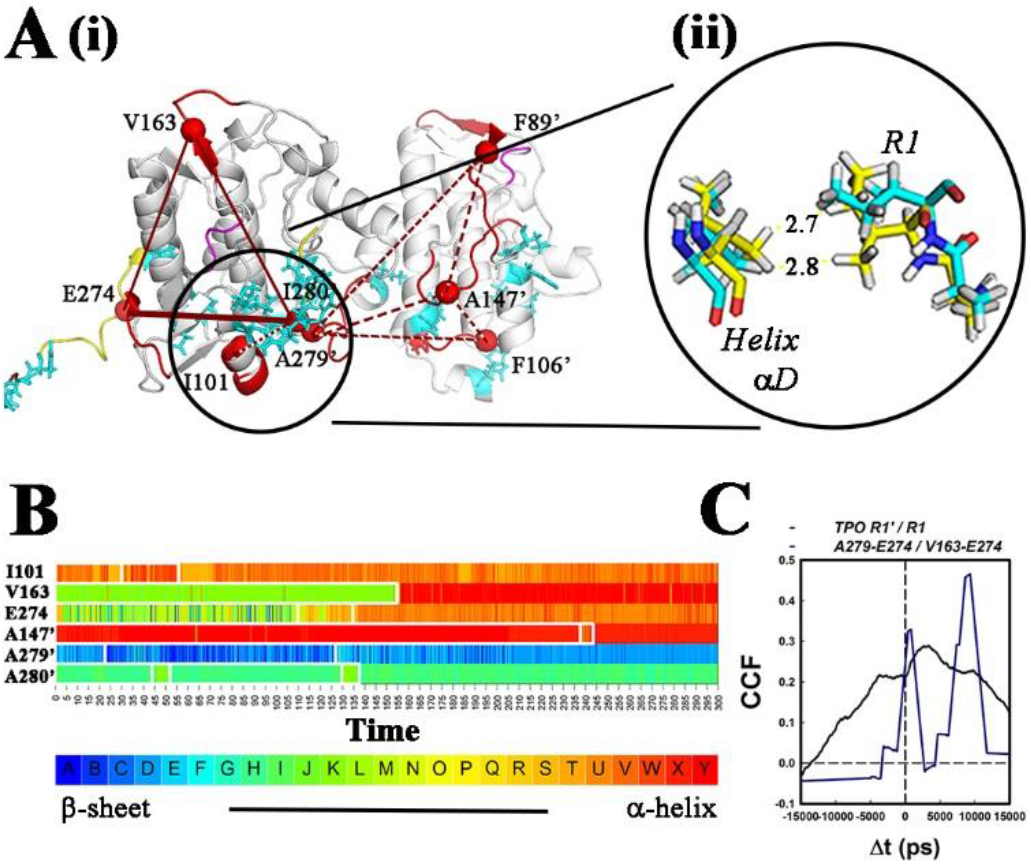
Temporal couplings of the R1-R1 signal network. **A. (i)** Parallel signal pathways formed by the dominant, long-range couplings in the C-lobe. The hydrophobic patch (circled) formed by helix αD and captured R1 orchestrates the couplings. Two pathways (1 direct (thick red line), 1 indirect (thin red lines) connect the patch and the free R1. The donor C-lobe also links to the patch (dotted red lines). R1 segments (yellow) and DFG motifs (purple) are highlighted. Cyan side-chains show residue positions where mutations cause learning disabilities in humans. **(ii)** atomic (stick) representation of the hydrophobic patch in the major configurations derived from cluster analysis of the T (cyan) and TPO (yellow) forms. **B.** Times series from one replica run of structural transitions within the signal pathway fragments (name = first residue). Colors denote SA letters as in B. **C.** CCF couplings between R1 segments in the phosphorylated (TPO) dimer (Dτ ~ 2.5 ns. τ_1/2_ ~ 10 ns) (black line). CCF of information flow to the free R1 from the adjacent C-lobe fragment correlated with the bound R1 fragment that forms part of the hydrophobic cluster.

Time series of the local, SA-encoded structural transitions for residue fragments that constitute this phosphorylation-dependent network are plotted for one replica run (Figure 8B). The most frequent transitions were short (< 10 ns) and sampled a restricted range of dihedral angles. Dihedral angle jumps scale with the separation between SA letters. Large jumps were rare but persisted for longer times (> 100 ns) when they occurred. Two examples are shown. The large jump for fragment V163 loop to α-helix is around the middle (~150 ns) of the time series. There is a smaller jump for fragment E274 between loop (β-sheet to α-helix) configurations at a similar time.

The cross-correlation (CCF) between the nMI time-series for the long-range (> 12 angstrom) couplings in the direct (A279-E-274) and indirect (V163-E74) R1-R1 pathways in the phosphorylated (TPO286) dimer network showed two peaks consistent with the short duration of fragment structural transitions (1-2 ns τ_1/2_’s) (Figure 8C). The first peak (Δt < 1 ns) presumably represents couplings associated with the direct pathway. The second peak (Δt ~ 10 ns) represents the lag associated with the first stage of the indirect, two-stage pathway. The CCF for the EED fluctuations of the two R1 segments in the phosphorylated dimer reported a weak correlation (amplitude = 0.35, correlation time τ = 15 ns). The correlation was not significant for the non-phosphorylated dimer.

## Discussion

Our findings indicate that the captured R1 freezes collective motions of the receiver subunit C-lobe to orchestrate allosteric coupling based on helix melting. Secondary structure fluctuations revealed α-helical character of free R1 compatible with ESR evidence (13) and showed that the captured RI transitions to an extended conformation energized by binding site solvation. The strained R1 triggered long-range couplings that propagated to the receiver R1. R1 is similar to α-helical linkers in the bacterial flagellar basal body that also respond to strain (39). Such “chameleon” helices may play essential roles in propagating conformational states across multiple subunits. While initial R1 capture has an enthalpic contribution, subsequent signal propagation is entropic as found for many systems (40).

The allosteric network (1) has a global architecture based upon a generic monomer KD network; (2) has retained nucleotide and substrate binding site couplings important for activation; (3) is strengthened by T286 phosphorylation and (4) incorporates mutated residue positions implicated in human disabilities as well as possible therapeutic targets. We discuss these network properties below.

### 1. Global architecture

Dynamic couplings consider both population shifts and propagation timescales of conformational ensembles (41). Thus, they provide a more refined readout for mechanistic analysis than the architecture of the protein fold (42). The auto-inhibited CaMKII form is completely inactive since the ATP binding site is not in an optimal conformation and the substrate binding site is occluded by the regulatory segment. The multiple community membership of the DFG and helix αD sites drove global adaptations of the network architecture coupling nucleotide and substrate binding. Reduced coupling between the communities, most marked for the coiled-coil dimer (BDW) structure, characterized the inactivated state. R1 capture results in reduced flexibility of the receiver C-lobe core; reported by RMSF, PCA, cluster and community network analysis.

The monomer network is co-opted in the chained dimer to generate a network that spans both KDs. A hydrophobic patch between the R1 central residues and helix αD orchestrated R1_donor_-> R1_receiver_ signal relays; most dominantly from the patch directly to R1_receiver_ N-and indirectly via the receiver DFG motif. Interfacial interactions linked the donor DFG motif to the patch. The prominent fragment couplings lasted a few to a hundred nanoseconds. Most were confined to short (~ 10 ns) transitions over a small conformational range. Large conformational transitions were infrequent but persisted for longer (>100 ns) times. Transitions between phosphorylated and dephosphorylated conformations completed within hundred nanoseconds. The temporal behaviour is consistent with a complex free-energy landscape with multiple pathways (43). We suggest the reduced flexibility of the receiver KD C-lobe upon R1 capture will influence the displaced receiver R1 to explore and bind to adjacent “open” subunits with floppy cores rather than re-associate with its own frozen, occupied core. The outcome is a self-reinforcing network that can be serially transmitted across the ring of KD C-lobes.

### 2. Activation dynamics

Our survey of CaMKII KD monomer structures revealed the nucleotide binding DFG motif and the B-site αD helix as conserved, coupled network nodes. The DFG motif is OUT when the KD is bound to the AD hub; an interaction analogous to the docking interaction of the PKA catalytic subunit with its regulatory subunit (7). There were two distinct DFG IN orientations, one associated with activated and the other with inactivated states, as in Aurora kinase (AurA) (44). The DFG motif is IN in auto-inhibited KD’s not bound to the hub. Thus, DFG motif orientation may report on compact and extended inactive holoenzyme states that have been visualized by EM cryo-tomography (45). In contrast, helix αD is OUT in all auto-inhibited, but IN for activated open and substrate-bound structures.

Our results add to data arguing that while the DFG motif is an important determinant of ATP binding its mobilization follows different strategies during activation of EPKs. In AurA, activation by the effector TpX2 involves an equilibrium population shift from the OUT to IN state (46); but activation by phosphorylation triggers a switch between auto-inhibited and activated IN states (44). Multiple strategies for kinase auto-inhibition have also been identified, for example, in the ZAP-70 tyrosine kinase (47).

Cooperativity in BCR-ABL, a tyrosine kinase identified as hallmark for myeloid leukemia can be either negative (48, 49) or positive (26) depending on the coupling between the nucleotide and substrate binding sites. In CaMKII, as reported here, the nucleotide and substrate binding modules are again recruited, but for formation of trans-subunit couplings. Comparative bioinformatics reveal common substrate binding site interactions between CaMKII and phosphorylase kinase (50), but the phosphorylase holoenzyme is constitutively active and its complexity limits study of nucleotide-substrate coupling.

### 3. T286 phosphorylation

T286 trans-phosphorylation has short and long-term consequences for LTP. In the short-term, second time scale TPO286 increases CaMKII affinity for calmodulin and decreased affinity for actin, triggering dissociation from the cytoskeleton and sequestration at the PSD in dendritic spines. It facilitates optimal Ca^2+^/ CaM binding, increasing its affinity for R2 more than 1000-fold and enables frequency-dependent activation (29, 51). It also alters the affinity of some substrates for CaMKII, thus modifying substrate selection (52, 53). *In vitro*, activation triggers subunit exchange over minutes to propagate the activated state to previously inactive holoenzymes and prolong the lifetime of an activated holoenzyme population (54). Synaptic localization of CaMKII holoenzymes triggered by millisecond calcium transients (55) persists over an hour or more (15), implying the activated CaMKII transitions to a long-lived state.

The MD simulations of T286 and TPO286 modified CaMKII dimers resolved changes triggered by primary site (T286) phosphorylation. T286 phosphorylation strengthened the coupling between chained KDs, with pK shifts of buried residues to more neutral values and surface, interfacial residues to more polar values, accompanied by compaction of the hydrophobic patch. The long-range allosteric network could regulate cytoskeletal interactions, calmodulin trapping and kinase cooperativity; all factors that would in turn modulate frequency tuning - a key parameter determining CaMKII physiology.

The dissociated holoenzymes must remain activated during diffusion to the PSD, so as to not rebind to the cytoskeleton. Similarly, during subunit exchange, the activated dimer must last longer than the time for encounter with another holoenzyme. Diffusion and encounter times are on the order of milliseconds based on known spine volumes (~ 1 μm^3^) and spine CaMKII concentrations (> 0.1 M). The increased Ca^2+^/ CaM binding affinity of the phosphorylated dimer extends the window for R1 capture beyond the transient calcium pulse (< 20 seconds). The greater stability of the phosphorylated versus non-phosphorylated chained dimer predicted here provides additional discrimination to prolong the activated state after Ca^2+^/ CaM dissociation. The nanosecond coupling dynamics accommodate the rapid shifts in calmodulin and actin affinity. We speculate the nanosecond fluctuations would be low-pass filtered in the dodecameric holoenzyme to freeze the activated cores over millisecond times for diffusion to the PSD or subunit exchange. Conformational spread can achieve long-lived (seconds) configurations in multi-subunit ring assemblies such as the rotation states of the bacterial flagellar motor (56).

The self-reinforcing R1 signal relays are likely to facilitate conformational spread. Structural elements that regulate the positive CaMKII kinase cooperativity as with Ca^2+^/ CaM concentration have been identified. Reported Hill-coefficients *(H)* for substrate phosphorylation varied from 4.3 to 1.1 with Ca^2+^/ CaM concentration accompanied by shifts in half-maximal dose. They are downshifted by inhibitor peptides (H 4.3 → 1.7), hydrophobic patch residue mutations (3.0 -> 1.5) and native / decoy KD mixtures (1.5 → 0.9) and correlate inversely with linker length (H =4.3 → 1.7) in the *C. elegans* holoenzyme (8). *H* is reduced (H 2.1 → 1.1) by residue mutations in the AD hub interface docking the KD DFG motif in the human holoenzyme (7). The inhibitor peptide and hydrophobic patch modulations of *H* values agree qualitatively with the main features of the dynamic network we have characterized. The linker length and AD mutations merit study of their role in Ca^2+^/CaM activation dynamics. Two distinct modes for conformational coupling have been proposed; lateral spread of the activated conformation across the holoenzyme subunits (8) or transverse paired dimers (9, 32). The latter may also mediate activation-triggered subunit exchange, driven by a strained, central AD hub (57) (**Supporting Information Figure S3**).

### 4. Physiological implications

The behavioural, histological and cellular phenotypes of the human mutations have been documented (18, 19), but these phenotypes are too downstream to decipher their molecular basis. This study shows that the mutated residue positions are distributed along the allosteric signal relays generated by R1 capture and strengthened by T286 phosphorylation to provide, for the first time, a molecular rationale. Namely that the mutations principally act to disrupt the allosteric network rather than weaken substrate or nucleotide binding per se. Disruption of the allosteric network could have multiple outcomes as discussed above to produce gain or loss of function with diverse pathophysiological consequences.

Importantly, this study opens possible avenues for therapeutic treatment. MD simulations have proved useful for determination of the efficacy of peptide inhibitors to ATP-binding pocket residue mutations (58). The design of allosteric inhibitors is more challenging as it requires evaluation of the conformational plasticity of the protein assembly but promises greater specificity. We have identified protein sectors based on community analysis that may disrupt conformational coupling without altering calmodulin, nucleotide or substrate binding. Their predicted role as signal relays rather than binding determinants can be tested by mutant screens. An excellent example in the literature of such a mutation is PKA Y204A that disrupts coupling between nucleotide and substrate binding without affecting binding affinities (59).

In conclusion, our simulations make the case that the conformational dynamics of chained dimers have advantageous properties for subunit exchange and holoenzyme activation. Network analysis revealed the centrality of the coupled A (DFG motif) and B (helix αD) sites in the R1 relay to suggest how substrate affinity is modulated by nucleotide occupancy and how both influence cooperativity. It detailed how T286 phosphorylation strengthened conformational coupling initiated by R1 capture. Finally, community graphs identified targets for rational design of allosteric inhibitors (22). Future work will build on these advances to reconcile the dynamic architecture of the kinase with measurements of its cooperativity.

## Materials & Methods

### 1. Phylogenetics

CaMKII sequences were retrieved from Uniprot (60). MUSCLE was used for multiple sequence alignment (MSA). Secondary structure predictions were made with PsiPred. The MSA was manually curated in Jalview. The phylogenetic tree was constructed with Fast-Tree 2.19 and displayed with Fig-Tree 4.3 (*http://tree.bio.ed.ac.uk/software/figtree/*).

### 2. Structure Preparation

The human δ KD structure alone (PDB ID: 2VN9) and with calmodulin (2WEL)(9), complete subunit from the human holoenzyme (3SOA (7)); the *C. elegans* KD structures alone (2BDW (23), 3KK8 (8)) and with CaMKIINtide (3KL8 (8)) and the open form of protein kinase A (1CMK (61)) were downloaded from Protein Data Bank. Missing atoms were added in Swiss-Pdb viewer (*www.expasy.org/spdbv*); missing loop segments with Modeller (https:/salilab.org/modeller). Mutant substitutions were made in Pymol (http://pymol.org) then energy minimized in Modeller.

### 3. tCONCOORD

Parameters for tCONCOORD runs were as detailed earlier (62). tCONCOORD utilizes a set of distance constraints, based on the statistics of residue interactions in a crystal structure library (25, 63), to generate conformational ensembles from an initial structure without inclusion of solvent. Sets of 16^4^ = 65,536 equilibrium conformations with full atom detail were generated for each structure. The overlap between ensemble subsets was > 99% when subset size was < ½ of this value.

### 4. Molecular Dynamics

A set of 3 replicas of 300ns were generated for *E. coli* 3KK8 structure and its 286T equivalent using Gromacs 2016.2 with Amber ff99sb*-ILDNP force-field (64). Each system was first solvated in an octahedral box with TIP3P water molecules with a minimal distance between protein and box boundaries of 12 Å. The box was then neutralized with Na+ ions. Solvation and ion addition were performed with the GROMACS preparation tools. A multistage equilibration protocol, modified from (65), was applied to all simulations to remove unfavourable contacts and provide a reliable starting point for the production runs including: steepest descent and conjugate gradient energy minimisation with positional restraints (2000 kJ mol^-1^ nM^-2^) on protein atoms followed by a series of NVT MD simulations to progressively heat up the system to 300 K and remove the positional restraints with a finally NPT simulation for 250 ps with restraints lowered to 250 kJ mol^-1^ nM^-2^. All the restraints were removed for the production runs at 300 K. In the NVT simulations temperature was controlled by the Berendsen thermostat, while in the NPT simulations the V-rescale thermostat (66) was used and pressure was set to 1 bar with the Parrinello-Rahman barostat. A set of 3 replicas with time step 2.0 fs with constraints on all the bonds were used. The particle mesh Ewald method was used to treat the long-range electrostatic interactions with the cut-off distances set at 12 Å. The threonine phosphate (TPO286) was changed to threonine (T286) after equilibration to generate the non-phosphorylated form.

### 5. Energetics

Contact residues, surfaces and energies were extracted from the PDB files with the sub-routines (*ncont, pisa*) available in CCP4 version 7 (*http://www.ccp4.ac.uk/*). Continuum electrostatics were computed with the Poisson Boltzmann solver (*http://www.poissonboltzmann.org/*)(67). Comparison with experimental B-factors and geometrical analyses were performed with GROMACS version 4.5.7 (*http://www.gromacs.org/*).

### 6. Structural alphabet

The mutual information *I*(*X; Y*) between two variables(*X*) and (*Y*) is

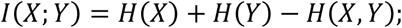

where *H*(*X, Y*) is the joint probability distribution;

The normalized mutual information, *nMI*(*X; Y*) = (*I*(*X;Y*) – *ε*(*X; Y*))/(*H*(*X, Y*)); where *ε*(*X; Y*) is the expected, finite-size error;

The *nMI* couplings are detected as correlated changes in fragment dynamics. In methods based on the dynamics cross-correlation matrix (68), the correlation is calculated from position fluctuations of the residue C^α^ atoms. In the MutInf method (69), correlations are measured in terms of mutual Information between discretized torsional angles. The residue-based approaches do not directly read-out couplings between structural motifs, e.g. secondary structures. The SA (70), is a set of recurring four residue fragments encoding structural motifs derived from PDB structures. There is no need for discretisation and / or optimisation of parameters as the fragment set is pre-calculated.

### 7. Principal Component Analysis

Collective motions were identified by PCA of the conformational ensembles. PCs were generated by diagonalization of the co-variance matrix of C^α^ positions. For tCONCOORD the motions have no time-scale, but comparison with MD trajectories was consistent with the notion that collective motions represented by the first few PCs are “slow” relative to smaller amplitude motions recorded by the later PCs.

### 8. Network Analysis

Statistically significant correlations between columns were identified with GSATools (33) and recorded as a correlation matrix. The correlation matrix was used to generate a network model; with the residues as nodes and the correlations as edges. The contribution of a node to the network was estimated by the eigenvector centrality, E, calculated directly from the correlation matrix:

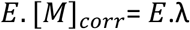

where [*M*]_*corr*_ is the correlation matrix and λ is the eigenvalue

The Girvan-Newman algorithm (34) was used to identify community structure. Then the network was collapsed into a simplified graph with one node per community, where the node size is proportional to the number of residues. Edge weights represent the number of nMI couplings between communities (35). Community analysis of correlation networks identifies relatively independent communities that behave as semi-rigid bodies. Graphs were constructed with the igraph library (71) in R (*https://cran.r-project.org/web/packages/igraph/*) and visualized in Cytoscape (*http://www.cytoscape.org/*).

### 9. Quantification and Statistical Analysis

Typical size of a tCONCOORD ensemble was 65,536 conformations (256^2^). Three MD replicate runs for the two (phosphorylated, dephosphorylated) conformations. Ensemble conformations and MD runs were averaged for computation of the nMI between fragment positions. Significance limits were set in GSATools. **Supporting Information Table S1** lists the web databases and software servers used in this study.

## Acknowledgments

AP was supported by Brunel BRIEF Award. H.S was supported by National Institutes of Health grant RO1 GM101277. SK had support from the Royal Society collaborative exchange grant U1175.70592 and seed funds from the Molecular Biology Consortium. Computation for the work described in this paper was supported by the Crick Data Analysis and Management Platform (CAMP), provided by the Francis Crick Institute.

## Author contributions

Conceptualization: AP and S.K. Methodology: AP and SK, Software: AP, Validation: SK, Writing - First Draft: SK. Writing -Review & Editing: AP, HS and SK, Visualization: AP, HS and SK, Supervision / Project Administration: SK. Funding Acquisition: AP, HS and SK.

## Supporting Information

Appendix S1: Table & Figures (PDF)

Figure S1: CaMKII KD Conservation

Figure S2: PKA and CaMKII have Homologous Structure and Dynamics.

Figure S3: Conformational coupling in the CaMKII holoenzyme.

Table S1: Web Resources.

Movie S1-S2. PCA of the 3KK8 monomer and dimer (AVI). The three major communities are colour-coded as in Figure 3A. TPO286 (red spheres). The filtered PC, PC1 and PC2 motions are shown. Related to Figure 4A.

Movie S3-S4: Filtered PC trajectory of the free and captured R1 (AVI). Three replica MD runs of the non-phosphorylated 3KK8 dimer sampled at 600 ps. T286 (yellow sphere). Residues are color-coded according to type (white = hydrophobic, red = acidic, blue = basic). Related to Figure 5A.

Movie S5: Network architecture of the phosphorylated 3KK8 dimer (AVI). Related to Figure 7C.

Movie S6: Surface profile and spatial relationship of intermediate relay community residues with the proline cluster targeted by human mutations and TPO286 (AVI). Related to Figure 7G

